# 2D:4D and spatial abilities: From rats to humans

**DOI:** 10.1101/193342

**Authors:** N Müller, S Campbell, M Nonaka, TM Rost, G Pipa, BN Konrad, A Steiger, M Czisch, G Fernández, M Dresler, L Genzel

## Abstract

Variance in spatial abilities are thought to be determined by in utero levels of testosterone and oestrogen, measurable in adults by the length ratio of the 2^nd^ and 4^th^ digit (2D:4D). We confirmed the relationship between 2D:4D and spatial performance using rats in two different tasks (paired-associate task and watermaze) and replicated this in humans. We further clarified anatomical and functional brain correlates of the association between 2D:4D and spatial performance in humans.

---

Many factors can influence performance on memory tasks and induce inherent between subject variability. For example, when investigating memory consolidation via spatial tasks, general spatial abilities will affect how well subjects perform. General spatial ability and preferred spatial strategy is a sexual dimorphic trait, which is thought to be influenced in utero via the testosterone/oestrogen hormone ratio [1]. This prenatal hormonal ratio does not correlate with sex hormone levels in the adult, making it very difficult to directly measure in memory experiments with adult subjects. Interestingly, this same prenatal hormone ratio also affects the growth of the 4^th^ digit with higher hormone ratios inducing longer 4^th^ digits in comparison to the 2^nd^ digit [2]. More specifically, androgen receptor (AR) and estrogen receptor α (ER-α) activity is higher in digit 4 than in digit 2. Inactivation of AR or activation of ER-α decreases growth of digit 4, which causes a higher 2D:4D ratio; whereas inactivation of ER-α or activation of AR increases growth of digit 4, which leads to a lower 2D:4D ratio. Intriguingly, several genes identified by Zheng et al to be responsible for the 2D:4D ratio also have roles in development of the brain [3]. Thus the ratio between these digits (2D:4D) is an indicator for the hormonal ratio in utero, with lower finger ratios indicating higher testosterone to oestrogen.

In a study aimed at investigating memory consolidation effects across different tasks in the same male animals, we noticed early in training while discussing the current state of performance that we had “smart” and “not so smart” rats across different tasks. And in fact, the correlation between probe trial performance in the Delayed-Match-to-Place version of the water maze, with daily switching escape locations [4], and in a flavour-location association task [5] in an open field environment (arena) was significant (n=20, r=0.53, p=0.015; fig 1A; see supplemental materials), suggesting some factor general spatial ability confounded the performance in both tasks. As a negative correlation between the right (but not left) 2D:4D and spatial performance in the watermaze has been demonstrated in humans [6], we dissected our rat’s front paws post-mortem and measured their 2D:4D. And in fact, the right 2D:4D correlated negatively with the performance in both tasks (2D:4D_arena: r=-0.46, p=0.044, 2D:4D_watermaze: r=-0.50, p=0.024; fig 1B,C). Similar to previous human results, there was no such correlation present using the left 2D:4D. Thus we are, to our knowledge, the first to translate the prenatal programming effect of sex hormones on cognitive domains to rats, further emphasizing that cognitive sexual dimorphic traits are not solely caused by societal influence but have an inherently biological component.

**Figure 1:**
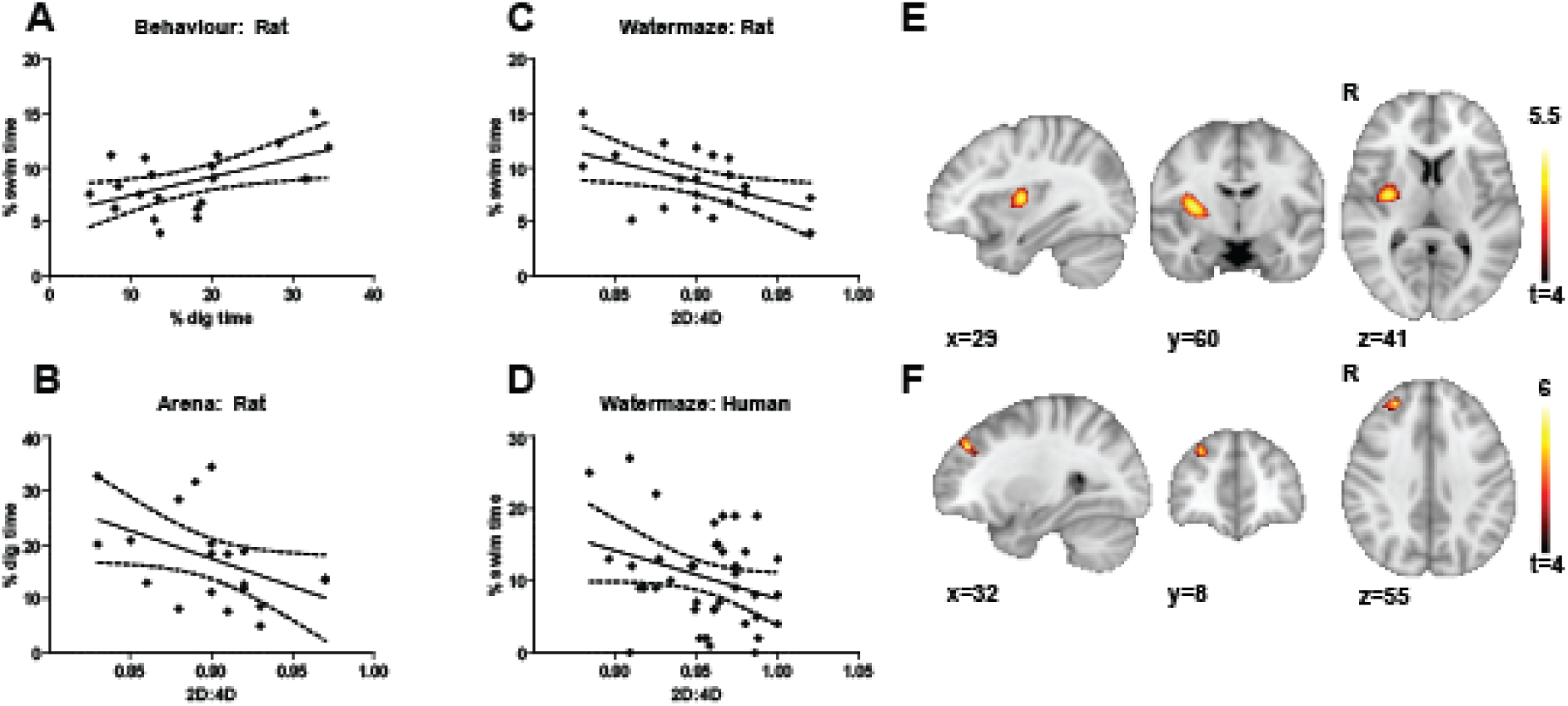
2D:4D, Behaviour and Neural Correlate. A. Shown is the performance in the water maze (y-axis) and arena task (x-axis) with significant regression line and 95%CI (F_1,18_=7.2, p=0.015). The regression analysis for both performance in the arena task and watermaze with 2D:4D was significant (B, arena F_1,18_=4.7, p=0.044; C, watermaze F_1,18_=6.1, p=0.024). Also in humans the performance in the watermaze showed a relationship with 2D:4D (D; F_1,38_=3.9, p=0.056). E. Anatomical analysis (VBM) showed a significant cluster in the right caudate that was positively associated with the 2D:4D index (p_FWE-whole brain_<0.05). F. Using the caudate cluster from A. as a seed, we could show that functional connectivity during rest between that cluster and a right frontal cortex region predicted the immediate water maze performance (p_FWE-whole brain_<0.01).

To investigate the neural basis of the association between 2D:4D and spatial ability, we turned to human subjects. We replicated our finding in rats that performance in the watermaze correlated with 2D:4D in a similar all male, human sample (n=40) using a virtual reality version of the watermaze. Performance in a probe trial after four learning trials correlated significantly with the 2D:4D index (r=- 0.30, p=0.028 one-sided; fig 1D). Further, we acquired structural as well as functional resting-state scans using MRI (see supplemental materials). We found a cluster including the right caudate nucleus and insula that showed a positive correlation with 2D:4D in the structural analysis (VBM, fig 1E); an area previously already identified as sexual dimorphic [7]. Interestingly, the positive correlation indicated that smaller 2D:4D (reflecting higher testosterone levels in utero) was associated to smaller size of this cluster. Further, including this anatomical cluster as a region of interest in the functional resting state analysis together with immediate watermaze performance as regressor, we found a significant cluster in the prefrontal cortex (p_FWE_whole brain_<0.01); thus the higher the resting state functional connectivity of the cluster found in the anatomical 2D:4D analysis with the prefrontal cortex cluster, the better the subjects performed in the watermaze, most likely indicating a more efficient spatial memory network (fig 1F).

With this study, we are the first to confirm the association between fetal sex hormones measured indirectly via 2D:4D and cognitive abilities in rats, reported previously in humans [6,8]. The testosterone/oestrogen balance in utero was shown previously to causally influence 2D:4D growth in mice via genes also present during development in the brain [2,3]. The testosterone levels in utero seem to be determined by the mother’s hormone production levels as well as testosterone produced by the fetus itself [9]. As shown here and previously in humans [6,8] these hormone levels seem to “preprogram” the brain to be either more adept in “male” tasks (e.g. spatial ability) or “female” tasks (e.g. verbal, recognition and fine motor skills; not tested here) by influencing brain maturation (i.e. anatomical and functional effects seen in fig 1 E,F).

The implications of this study are twofold. Firstly, it provides evidence for the biological are not solely social basis of “male” and “female” memory and cognition [1] and emphasizes that sex matters in neuroscience. Sex and sex hormones influence and can confound seemingly unrelated research and the 2D:4D is a possible marker to control for the influence of fetal hormonal influences. A second implication of this study is the confirmation of 2D:4D as cross-species biological marker for prenatal programming by sex hormones. Studies using 2D:4D to investigate prenatal programming effects of sex hormones on behavior and disease are increasing. For example, prostate and breast cancer were found to be associated with 2D:4D [10]. In summary, our study could show an association and neural correlate between spatial task performance and 2D:4D in rats and humans, providing further evidence for the biological nature of sex hormone effects on cognition [1]. These findings should be followed up with interventional approaches investigating how sex hormones affect brain development and subsequent spatial abilities.

## Acknowledgments

This study was supported the Branco Weiss, Society in Science Fellowship.

## Online Methods

### Rat Behavior

We trained 22 rats in two tasks: the watermaze and flavour-location paired-associates in an event arena. For the water maze the delayed-matched-to place version was selected[4] with the rats learning a new platform location within the watermaze (r=1m) every session (every ∼3 days) with 4 trials per session (15s inter-trial interval) starting each once from north, south, east and west (starting sequence and platform location counterbalanced). To test for the memory of the previous training day’s location the first trial of the day contained a probe trial using the Atlantis platform, which only became accessible after a 60 sec delay. The average swim time in the zone around the previous platform position (r=20cm, chance level 4%) of sessions 5-8 were used as outcome measure. The rats were additionally trained every 2^nd^ day in an event arena task with 6 flavour-location paired-associates[5]. A training session consisted of each one trial for every flavour-location associate, with a flavour cue given in one of four startboxes (located north, east, south, west) and the animal allowed to dig in 6 available sand wells for three additional pellets placed in the cued location (fixed for every flavour). After the first and then every six training sessions an unbaited probe trial (all sand wells were empty) was performed with %dig time measured for the cued sand well in comparison to the total dig time within the 120s probe trial (chance level 16.67%). We used probe trial 4 in the arena as outcome measure, which corresponded in time with sessions 5-8 in the watermaze. To test spatial abilities we used probe trial performance early in training with only individual good performance of some rats in each task.

After culling the animals the 2^nd^ and 4^th^ finger of the right paw was fixed in 4% formalin, dissected, measured (each digit x3 and then averaged) and the ratio calculated by an experimenter blind to the study design and behavioural performance. The paws of 2 animals were lost due to technical difficulties resulting in an n of 20.

### Human Behaviour

We tested 40 male participants in a virtual reality version of the watermaze task. Differing from the animal version the learning and testing was all done in a single session. Participants wore a head mounted display, a sensor for tracking their movements, and a joystick to navigate. Before the actual experiment, participants learnt to navigate first in an independent setting (unlimited time) and then had a short familiarization (60s) with the watermaze without the hidden platform. Afterwards the 4 learning trials started in which they had to find a hidden platform (7 units in diameter) in the watermaze (100 units in diameter). After each trial, participants had a 5 min. break during which they filled out questionnaires in order to prevent rehearsal. The fifth trial was a probe trial in which the platform was accessible after a 60 seconds delay. The swim time in the area around the platform (r=14 units) during these 60 seconds was used as outcome measure.

We obtained the 2D:4D index by using a generic copy machine and asking participants to apply slight pressure when putting the hand down on it. The printed out copies where then measured by 2 student assistants blind to the study design and behavioural performance and their results averaged.

### Human MRI

In a separate session imaging data were collected using a 3T (GE Discovery MR750) scanner with a 12-channel head coil. A standard localizer, coil calibration and a 3D T1-weighted anatomical scan (TR7.1 ms, TE 2.2 ms, slice thickness 1.3 mm, in-plane FOV 240 mm, 320×320x128 matrix, 12° flip angle) preceded fMRI data collection. Eight minutes of resting state fMRI with eyes closed were collected (EPI sequence, TR 2.5 s, TE 30 ms), covering the whole brain with 34 slices, using a 64×64 matrix with 3 mm slice thickness and 1 mm slice spacing, and a field of view of 240 x 240 mm2. The images were AC–PC aligned and acquired using an interleaved slice acquisition scheme.

We analysed structural brain scans with FSL-VBM [11] using an optimised VBM protocol [12]. Preprocessing included non-linear registration to a study-specific template and smoothing with an 8 mm isotropic Gaussian kernel. The regression model used included whole brain size, age, IQ, hours of video game experience as covariates and 2D:4D and the watermaze performance as regressors of interest. Statistical inference testing was done using a permutation test with 15.000 permutations. All results were thresholded at p<0.05 family-wise error correction using threshold free cluster enhancement implemented in FSL randomise.

To test whether they grey matter differences relate to functional connectivity differences we did a seed-based correlation analysis using resting state data. Preprocessing included motion correction, smoothing with a 6mm isotropic Gaussian kernel, slice time correction, bandpass filtering (0.01 to 0.1 Hz) and a nonlinear registration to MNI152 space. For every voxel outside the ROI obtained from the VBM analysis we calculated the average correlation with the ROI. These whole-brain correlation maps where then used in the identical regression analysis used in the VBM analysis.

